# 3.1Å structure of yeast RNA polymerase II elongation complex stalled at a cyclobutane pyrimidine dimer lesion solved using streptavidin affinity grids

**DOI:** 10.1101/585547

**Authors:** Indrajit Lahiri, Jun Xu, Bong Gyoon Han, Juntaek Oh, Dong Wang, Frank DiMaio, Andres E Leschziner

**Affiliations:** Department of Cellular and Molecular Medicine, University of California San Diego, La Jolla, CA, 92093, USA; Division of Pharmaceutical Sciences, Skaggs School of Pharmacy & Pharmaceutical Sciences, University of California San Diego, La Jolla, CA, 92093, USA; Molecular Biophysics and Integrated Bioimaging Division, Lawrence Berkeley National Laboratory, University of California, Berkeley, CA, 94720, USA; Department of Biochemistry, University of Washington, Seattle, WA, 98195, USA; Division of Biological Sciences, University of California San Diego, La Jolla, CA, 92093, USA

## Abstract

Despite significant advances in all aspects of single particle cryo-electron microscopy (cryo-EM), specimen preparation still remains a challenge. During sample preparation, macromolecules interact with the air-water interface, which often leads to detrimental effects such as denaturation or adoption of preferred orientations, ultimately hindering structure determination. Randomly biotinylating the protein of interest and then tethering it to a cryo-EM grid coated with two-dimensional crystals of streptavidin (acting as an affinity surface) can prevent the protein from interacting with the air-water interface. Recently, this approach was successfully used to solve a high-resolution structure of a test sample, a bacterial ribosome. However, the general applicability of this method to samples where interaction with the air-water interface is problematic remains to be determined. Here we report a 3.1Å structure of a RNA polymerase II elongation complex stalled at a cyclobutane pyrimidine dimer lesion (Pol II EC(CPD)) solved using streptavidin grids. Our previous attempt to solve this structure using conventional sample preparation methods resulted in a poor quality cryo-EM map due to Pol II EC(CPD)’s adopting a strong preferred orientation. Imaging the same sample on streptavidin grids led to a high-resolution structure with little anisotropy, showing that streptavidin affinity grids could be used as a general strategy to address challenges posed by interaction with the air-water interface.

## Introduction

Single particle cryo-electron microscopy (cryo-EM) is quickly becoming the technique of choice for obtaining the structures of biological macromolecules at atomic or near-atomic resolutions (Nogales and Scheres, 2015). In this method, the sample is applied to an EM grid and excess liquid is blotted away, leaving a thin film of buffer containing the macromolecules. The grid is then rapidly frozen in a cryogen such that the thin buffer film vitrifies into amorphous ice with the macromolecules embedded in it. A recent study has shown that, in most cases, instead of being uniformly distributed in the thin layer of vitrified buffer, the macromolecules become adsorbed to the air-water interface (Noble et al., 2018a) indicating that prior to vitrification the macromolecules interact with the air-water interface of the thin film. This interaction can have detrimental consequences, including complete or partial denaturation of the sample or the adoption of preferred orientations by the macromolecules, ultimately hindering structure determination (Glaeser and Han, 2017).

Currently, there are three major approaches to prevent or minimize the exposure of samples to the air-water interface: (a) Including small molecule additives in the solution to alter its properties (Bernecky et al., 2016; Kang et al., 2017); (b) minimizing the time between thin film formation and vitrification (Noble et al., 2018b); and (c) adding an affinity tag to the macromolecule to “tether” it to an affinity surface, keeping it away from the air-water interface (Han et al., 2012; Kelly et al., 2010; Sharma et al., 2013). While the first approach has been used successfully in many cases, its effectiveness is dependent on the nature of the sample, and its implementation requires extensive optimization. The second approach, speeding up the vitrification process, is very promising but it is a relatively new technique still under development (Dandey et al., 2018). The affinity-tagging approach has the potential for reproducibly preventing proteins from reaching the air-water interface and could easily be applied to a wide variety of macromolecules. Genetically-encoded affinity tags are limiting for this approach; placing the tag in a particular location restricts the orientations the sample can adopt on the affinity support, negatively affecting structure determination. This issue can be circumvented by randomly tagging the protein, such as with biotin. The randomly biotinylated protein can then be attached to a streptavidin monolayer affinity grid (Han et al., 2012). This approach was successfully used to solve a near-atomic resolution structure of a bacterial ribosome (Han et al., 2016). However, it remained to be determined whether streptavidin affinity grids could be a general strategy to solve high-resolution structures of samples where interaction with the air-water interface has been shown to be problematic.

We have used streptavidin grids to solve the structure of a randomly biotinylated yeast RNA polymerase II elongation complex stalled at a cyclobutane pyrimidine dimer (CPD) lesion (Pol II EC(CPD)) to 3.1Å. The same sample had displayed a strong orientation bias when applied to holey carbon grids (both with and without an additional carbon or graphene support), severely limiting the attainable resolution (Xu et al., 2017). To our knowledge, the present study is the first instance showing that streptavidin affinity grids can be used to overcome a known preferred orientation problem, enabling structure determination to high resolution.

## Methods

### Purification and biotinylation of Pol II Elongation Complex arrested at a CPD lesion (Pol II EC(CPD))

We purified *S. cerevisiae* 12-subunit Pol II and formed a CPD-arrested elongation complex as previously described (Xu et al., 2017). After forming the elongation complex (EC) we dialyzed it into buffer A (40mM HEPES (pH7.5), 5mM MgCl_2_, 40mM KCl, 5mM DTT). For biotinylation of Pol II EC(CPD), we added 25-fold excess of biotin-labeling reagent (ChromaLink Biotin, Trilink, USA) to the sample and incubated the reaction for 2h at room temperature. We stopped the reaction by adding Tris-Cl (pH 7.5) to a final concentration of 50mM. We removed excess unreacted biotin by dialyzing the sample overnight against buffer A. To test the crosslinking efficiency, we incubated the Pol II EC(CPD) (with and without biotin crosslinking) with 2μl streptavidin magnetic beads (NEB, USA) for 30 min at room temperature. The beads were then pelleted using a magnetic stand and the supernatant, containing Pol II EC(CPD) not bound to the streptavidin beads, was run on a 4.5% native polyacrylamide gel in TBE buffer (pH 8.0) with 2mM MgCl_2_ for 2.5h at 4 °C and stained with Gel-red. We determined the extent of biotinylation based on the relative intensities of the Pol II EC(CPD) bands between the experiment and no-bead control. Based on these calculations ~ 30% of Pol II EC(CPD) was biotinylated. After biotinylation the sample was dialyzed in buffer A to remove free biotin reagent.

### Electron microscopy sample preparation and imaging

We applied a streptavidin crystal monolayer to Quantifoil R2/2 grids (gold grids with carbon foil) as previously described(Han et al., 2016). The streptavidin layer was protected by an additional layer of trehalose and we stored the grids in a desiccator. The protective trehalose coating was removed by washing the grids with rehydration buffer (50mM HEPES (pH 7.5), 150mM KCl, 5mM EDTA) just prior to sample application. We applied 4μl of 300 nM Pol II EC(CPD) (where an estimated 30% of the sample is biotinylated) to the affinity grid and incubated it in a high humidity chamber for 10 minutes. Unbound sample was washed with Buffer A and then we used a Vitrobot (FEI) to blot away excess sample and plunge freeze the grid in liquid ethane. The blotting was performed as previously described (Han et al., 2016). The grids were stored in liquid nitrogen until imaged.

For negative stain EM we used a JEM-1200EX (JEOL) TEM for imaging. The microscope was operated at 120 kV with a nominal magnification of 32,000x corresponding to an object-level pixel size of 4.3Å. For cryo-EM we imaged the grids in a Talos Arctica (FEI) operated at 200□kV, equipped with a K2 Summit direct detector (Gatan). Automated data collection was performed using Leginon (Suloway et al., 2005). We recorded 1,800 movies in ‘counting mode’ at a dose rate of 11.6 electrons pixel^−1^ s^−1^ with a total exposure time of 6s sub-divided into 150ms frames, for a total of 40 frames. The images were recorded at a nominal magnification of 36,000x resulting in an object-level pixel size of 1.16Å. The defocus range of the data was −0.5μm to −3.5μm.

### Image processing

We aligned the movie frames using UCSF MotionCor2 (Zheng et al., 2017) as implemented in Relion 3.0 (Zivanov et al., 2018), using the dose-weighted frame alignment option. We removed the signal for the streptavidin lattice from the aligned micrographs using Fourier filtering as described before (Han et al., 2012). The scripts used for Fourier filtering are available at the accompanying Mendeley data site http://dx.doi.org/10.17632/k2g2p5z9x6.1#folder-601061c5-3fed-4902-a747-6b7a45fa01d2). The micrographs were then inspected and the unsuitable ones (having defects like broken carbon, crystalline ice or too thick or thin an ice layer) were discarded. We estimated CTF on the non-dose-weighted micrographs using GCTF version 1.06 (Zhang, 2016) as implemented in Appion (Lander et al., 2009) and micrographs having a 0.5 confidence resolution for the CTF fit worse than 4.5Å (as determined in Appion) were excluded from further processing. Using this approach, 1,200 micrographs were kept for further processing. We selected particles from the micrographs using FindEM (Roseman, 2004) with projections of a previously reported map of a Pol II EC (EMDB 8737) (Xu et al., 2017) serving as references. Particle picking was performed within the framework of Appion.

We carried out all subsequent processing in Relion 3.0 installed in Amazon Web Services (Cianfrocco et al., 2018; Cianfrocco and Leschziner, 2015). Two-dimensional classification was performed to identify and remove ‘bad’ particles. Only those particles that contributed to ‘good’ 2D class averages were used for further processing. Following 2D classification, we performed a 3D classification using a Pol II EC model (PDB accession 1Y77) (Kettenberger et al., 2004) as reference. The reference was low pass filtered to 60Å prior to use. 61,654 particles contributing to the class showing good overall density and characteristic structural features of Pol II EC (for instance, the bridge helix) were selected for further processing. We performed the 2D and initial 3D classifications using binned data (3.9Å pixel^−1^) and all subsequent steps using unbinned images (1.16Å pixel^−1^). The classification and refinement scheme is shown in Figure S2. After the consensus refinement, we estimated CTF for each particle and beam tilt for each micrograph. We performed an additional round of refinement using the per-particle CTFs and the beam tilt parameters. The resolution of the cryo-EM map was estimated from Fourier Shell Correlation (FSC) curves calculated using the gold-standard procedure and the resolutions are reported according to the 0.143 cutoff criterion (Henderson et al., 2012; Scheres and Chen, 2012). FSC curves were corrected for the convolution effects of a soft mask applied to the half maps by high-resolution phase randomization (Chen et al., 2013). For display and analysis purposes, we sharpened the map with an automatically estimated negative B factor as implemented in the ‘post-process’ routine of Relion. We calculated 3D FSC using 3dfsc version 2.5 (Tan et al., 2017) using the half-maps and the mask used for the post-process routine in Relion. Unbending of the streptavidin crystal lattice in the micrographs was performed as described previously (Han et al., 2016) using the 2dx software package(Gipson et al., 2007). The first observed lattice spots (1,1 or −1,1) correspond to an approximate resolution of 57Å. We used Rosetta and Phenix (Adams et al., 2010; DiMaio et al., 2015; Song et al., 2013; Wang et al., 2016) to build the atomic model of Pol II EC(CPD) in our cryo-EM map following a previously described procedure (Xu et al., 2017). All figures and difference maps were generated using UCSF Chimera (Pettersen et al., 2004) and PyMOL (version 2.0 Schrödinger, LLC). The EM maps were segmented using Seggar (Pintilie et al., 2010) as implemented in UCSF Chimera. The electrostatic surface of Pol II was calculated using the APBS plugin (Baker et al., 2001; Dolinsky et al., 2004) in PyMOL.

### Estimation of solvent accessibility of amines

We estimated the solvent exposed surface area (SASA) of the amines using the LeGrand and Merz method (Le Grand and Merz, 1993) as implemented in Rosetta. Since the stalk region was disordered in our Pol II EC(CPD) structure, the corresponding atomic model does not include rpb4 and rpb7, the two Pol II subunits that form it. Therefore, to take into account the amino acids of rpb4 and rpb7 we used a previously reported atomic model of a Pol II elongation complex (Pol II EC) (PDB ID: 1Y77) (Kettenberger et al., 2004) for calculating SASA. Following sidechain rotamer optimization, solvent accessibility calculations were performed on all protein amine groups using a range of probe radii (1.0, 1.4, 1.8, 2.2, 3.0, and 4.0Å) to get 6 different accessibility values per target. We subsequently averaged these values to yield a single accessibility value. We performed this calculation only for the amine nitrogen and attached hydrogens (by considering only the surface points corresponding to these particular atoms). Averaging over a range of probe radii allows for softness in the calculation, making measurements less sensitive to small movements in amine groups or other nearby atoms, and a higher density of sampling near water radii values ensure those probe sizes dominate the average. We have implemented a sub-routine in Rosetta for this calculation (will be available from Rosetta 2019.10) and the instructions and scripts for executing this routine are provided in the accompanying Mendeley data site DOI: http://dx.doi.org/10.17632/k2g2p5z9x6.1#folder-f97fc421-396b-43c9-9445-9da401053780.

## Results and Discussion

### Biotinylation of an RNA Polymerase II elongation complex arrested at a DNA lesion

We first tested whether we could efficiently biotinylate an RNA Polymerase II (“Pol II”) Elongation Complex (“Pol II EC”) arrested at a cyclobutane pyrimidine dimer (“CPD”) DNA lesion (“Pol II EC(CPD)”). We incubated Pol II EC(CPD) with a 25-fold molar excess of the biotinylation reagent for 2 hours (Figure 1A). After terminating the reaction, we incubated the sample with streptavidin magnetic beads for 30 minutes. We removed the beads using a magnetic stand and analyzed the unbound fraction (Figure 1A) using native PAGE (Figure 1B). The unbound fraction represents the Pol II EC(CPD) that was not biotinylated. Based on the band intensities in the biotinylated and negative control (no biotinylation reagent added) samples we estimated that ~30% of the complexes were biotinylated (Figure 1B). Given this amount of biotinylation, we expect that the most abundant species in the biotinylated fraction will be Pol II EC(CPD) containing a single biotin moiety. We could obtain good particle density after incubating 4μl of 300nM Pol II EC(CPD) on the streptavidin grids for ~10 minutes (Figure1C, bottom panel). Given the extent of biotinylation in our reaction, this corresponds to 90nM of biotinylated Pol II EC(CPD). The high particle density was dependent on the presence of the streptavidin crystal; when we applied the same biotinylated sample to a continuous-carbon grid and the grid was subsequently washed to remove unbound complex, we detected very few particles in the micrographs (Figure 1C, top panel). The concentrating effect of the streptavidin grids is a major advantage of this approach.

**Figure 1:**
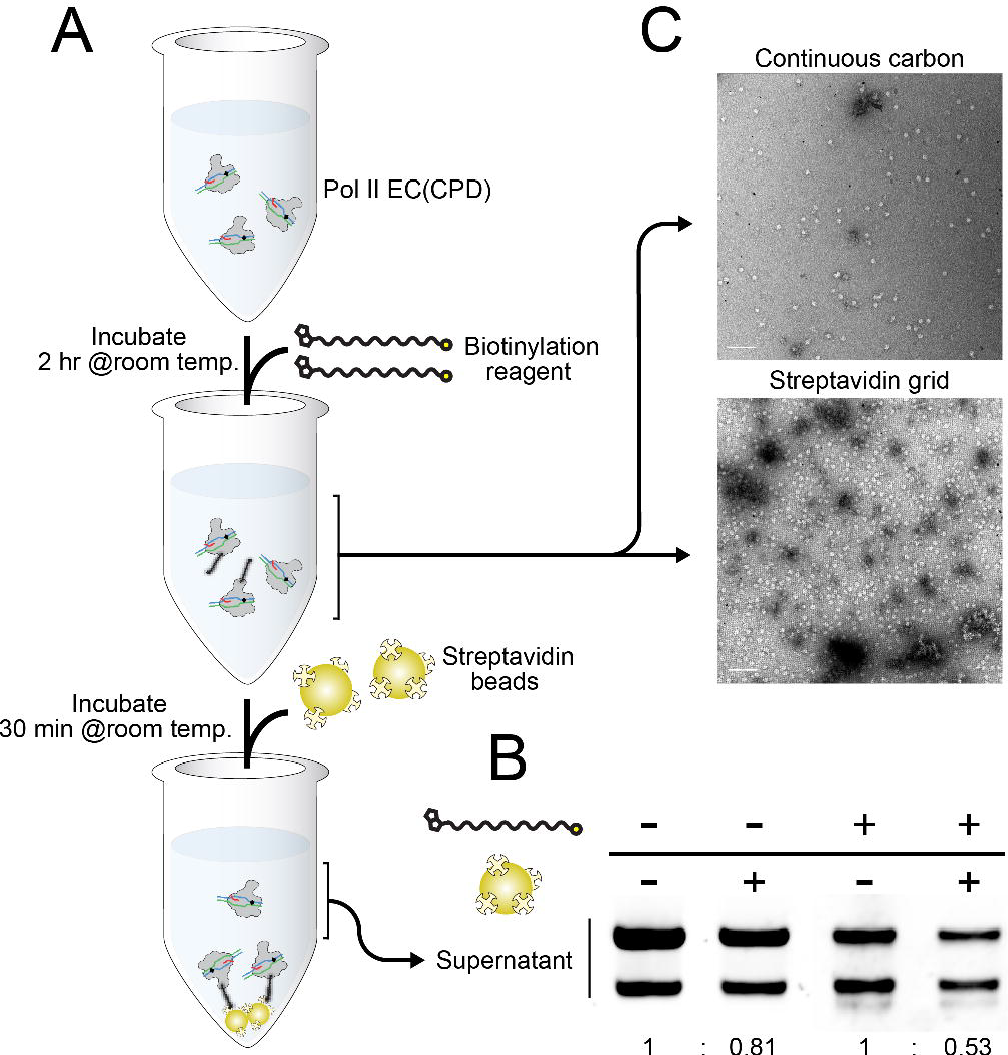
Biotinylation of an RNA Polymerase II Elongation Complex stalled at a cyclobutane pyrimidine dimer (CPD) lesion (Pol II EC(CPD)). **(A)** Schematic diagram of the assay used to determine the degree of biotinylation of Pol II EC(CPD). **(B)** Quantification of sample biotinylation. Samples were incubated with or without biotinylation reagent, and then with streptavidin magnetic beads (or without them as negative control). After pelleting the beads using a magnetic stand, supernatants were run on native PAGE gel. The ratios shown at the bottom indicate the streptavidin bead-dependent depletion of Pol II EC(CPD) in the absence (left) or presence (right) of biotinylation reagent. The two bands in each lane represent two different post-translationally modified states of Pol II EC(CPD) present in our sample (Sarker et al., 2005). **(C)** Negative stain EM micrographs of biotinylated Pol II EC(CPD) on continuous carbon (top) and streptavidin monolayer (bottom) grids.

### A 3.1Å structure of randomly biotinylated Pol II EC(CPD) on streptavidin monolayer grids

We prepared cryo-EM grids using a Vtrobot (FEI) with an additional manual wicking step as previously described(Han et al., 2016), and imaged them using a Talos Arctica (FEI). Using this approach, we could record images of Pol II EC(CPD) with good ice thickness showing good particle distribution over a background of uniform streptavidin crystal lattice (Figure 2A-C). We recorded a set of 1,800 movies. We found that more than 90% of the micrographs had a monolayer crystal of streptavidin, consistent with previous reports (Han et al., 2016). Discarding micrographs based on ice thickness (too thick, too thin or crystalline ice) and uniformity of the streptavidin crystal resulted in a set of 1,200 images that were further processed. The first step was removing the signal from the streptavidin crystal lattice using Fourier filtering (Figure 2D and E) followed by selecting particles using FindEM (Roseman, 2004). Two D and 3D classification (Figure S2) identified a homogeneous subset of 61,654 particles, which, after refinement, resulted in a 3.1Å map (based on the 0.143 cutoff of the gold-standard FSC curve) (Figures 3A and 4A). Although all 12 subunits of Pol II were present in our sample, the rpb4/7 stalk module was disordered in our EM map. The local resolution of the map showed that most of the complex was well resolved, with some parts, including the active site, having resolutions better than 3Å (Figure 3E). As has been seen in other Pol II EC structures (Bernecky et al., 2016; Liu et al., 2018; Xu et al., 2017) the flexible jaw domain showed the worst local resolution (Figure 3E). We could easily identify densities for many of the amino acid sidechains and nucleotides as well as for the CPD lesion (Figures 3B-D and G). To the best of our knowledge, the map presented here is the highest resolution cryo-EM structure of a Pol II transcription complex along with EMDB entries 0031 and 0036(Vos et al., 2018), both of which were obtained using data collected in a Titan Krios.

**Figure 2:**
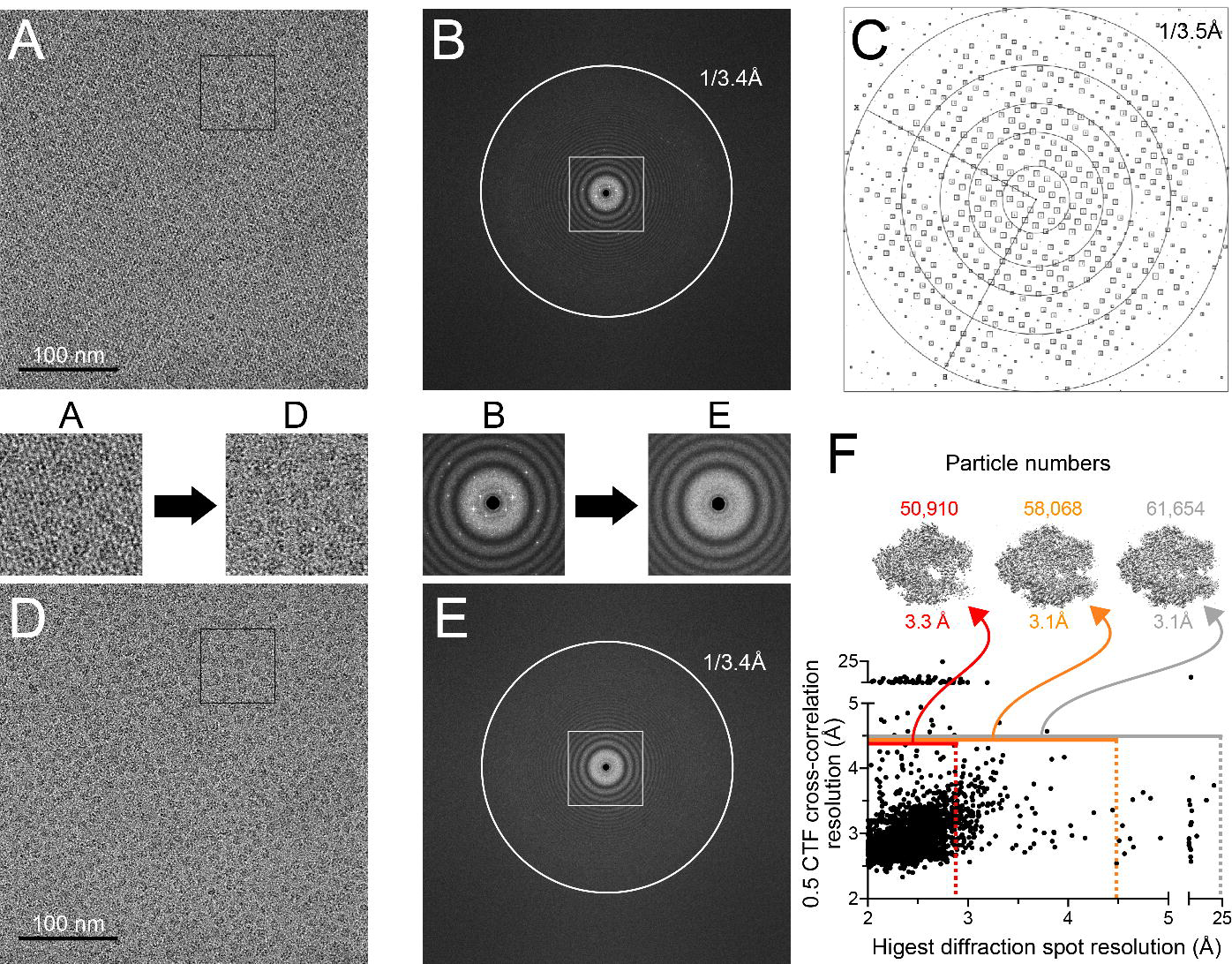
Processing of the streptavidin lattice in our images of Pol II EC(CPD) on streptavidin affinity support grids. **(A)** A representative micrograph showing Pol II EC(CPD) particles on a streptavidin crystal lattice. **(B)** FFT of the micrograph in (A) showing diffraction spots from the streptavidin crystal lattice. **(C)** Plot of diffraction spots of the streptavidin monolayer crystal from the micrograph shown in (A) after correcting the distortions in the 2D crystal (IQ plot) (Henderson et al., 1986). Each diffraction spot is shown as a square and the size of the square indicates the signal to noise ratio for that spot. **(D)** The same micrograph shown in (A) after removing the streptavidin lattice using Fourier filtering. **(E)** FFT of the micrograph in (D) showing the disappearance of the diffraction spots. Zoom-ins of the regions highlighted in (A) and (D) (black squares), and (B) and (E) (white square) are shown below (for A and B) or above (for D and E) the corresponding panels. **(F)** Plot of the resolution at which the cross-correlation between the calculated and observed CTFs is 0.5 as a function of the estimated resolution of the streptavidin crystal. The grey horizontal line shows the CTF resolution cutoff used for image processing (4.5Å). All the micrographs above the grey line were discarded. The grey (no cutoff), orange (4.5Å cutoff) and red (2.9 Å cutoff) vertical dashed lines show the diffraction spot resolution cutoffs used for different reconstruction trials. Micrographs to the left of the dashed lines (shown by horizontal lines of the corresponding colors) were used for the reconstructions. The resolutions of the maps from these trials along with the number of particles contributing to the reconstructions are shown above the graph. The map reported in this work was the one obtained without imposing a diffraction resolution cutoff.

**Figure 3:**
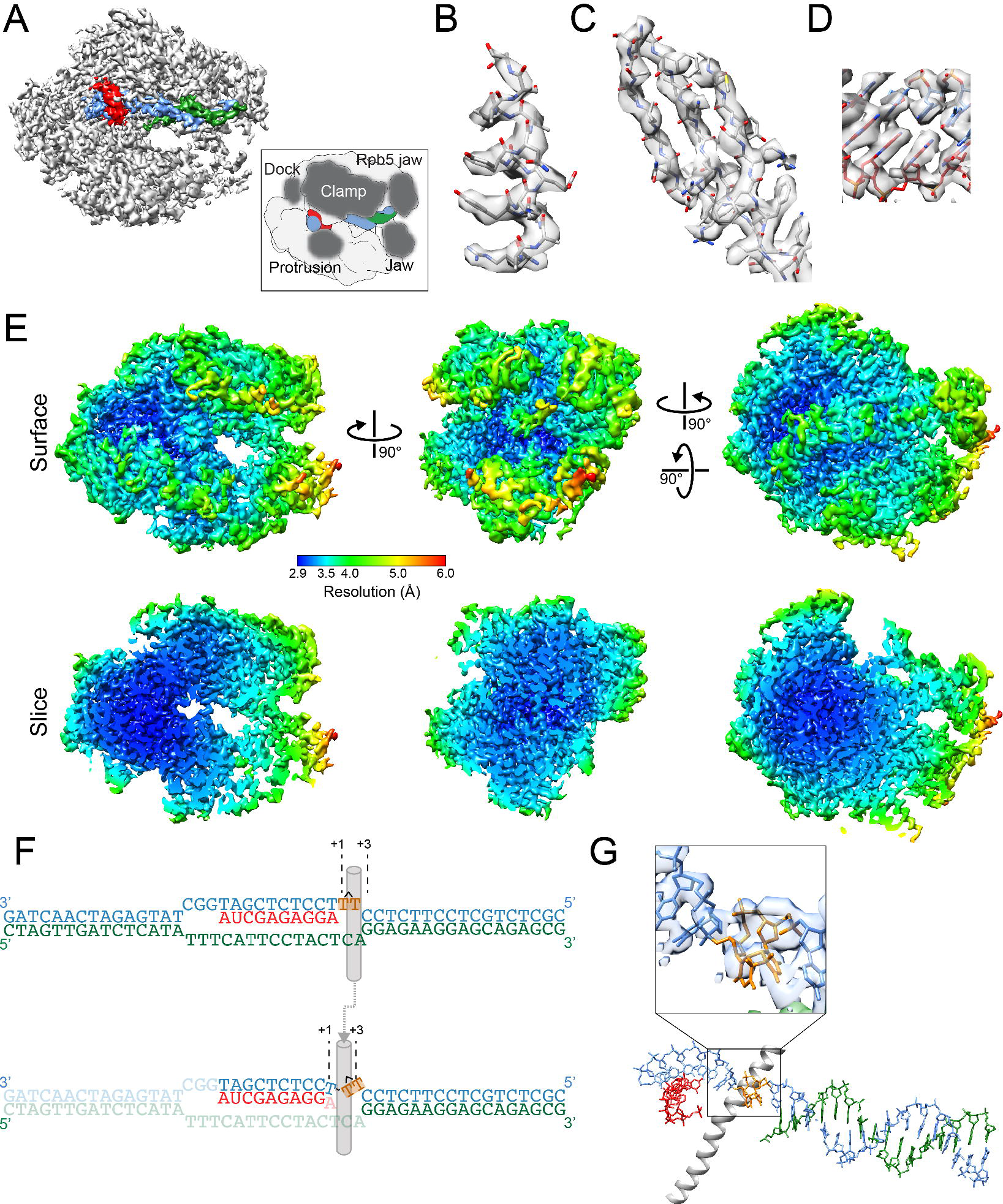
3.1Å cryo-EM map of a randomly biotinylated Pol II EC(CPD) obtained using monolayer streptavidin crystal affinity grids. **(A)** Locally-filtered Cryo-EM map of Pol II EC(CPD). The map is segmented into the following regions: Pol II (grey), template strand (TS) (blue), non-template strand (NTS) (green) and RNA strand (red). The inset highlights two domains of Pol II we refer to in the text on a cartoon representation of the structure. **(B-D)** Representative regions of the Pol II EC(CPD) map: an alpha helix (B), a beta sheet (C), and Watson-Crick base pairing between the TS and the nascent RNA (D). The map is shown in transparent grey with the atomic model fitted into the density. The color code for the atomic model is the same as in A. **(E)** Pol II EC(CPD) map colored according to local resolution. The top panel shows the surface views of the map in different orientations and the bottom panel shows the corresponding cross-sections to highlight the resolutions in the interior of the map. **(F)** Schematic representations of the expected (top) and observed (bottom) positions of the bases in the transcription scaffold. The grey cylinder represents the bridge helix. For the observed (bottom) scaffold the dash indicates that the CPD lesion (orange) is connected to the TS by a phosphodiester bond. Bases that are disordered in our map are shown in lighter color. The 5’ and 3’ ends of the TS (blue) and NTS (green) are indicated. **(G)** Top view of the atomic model of the transcription scaffold. The bridge helix is shown for reference. The inset shows the CPD lesion fitted to the segmented TS density.

It has been shown before that the quality of the streptavidin monolayer crystals could be used as a criterion for selecting particles that result in structures with improved resolution (Han et al., 2017). In our hands, this approach did not improve the map (Figure 2F). A possible reason might be the relatively small number of particles that were selected after 3D classification for the final reconstruction; additional selection based on the monolayer crystal quality further reduced the number of particles, likely offsetting the advantages provided by the selection. The results reported here show that even without considering the monolayer crystal quality, streptavidin affinity grids provide a robust approach to overcome the preferred orientation problem and solve structures of biological macromolecules to high resolution.

### Using streptavidin affinity grids increased the distribution of Pol II EC(CPD) views

The Pol II EC(CPD) map reported here has significantly higher resolution and lower anisotropy than our previously published structure of the same complex solved over open holes(EMDB 8885) (Xu et al., 2017). The current structure has an overall resolution of 3.1Å and a sphericity value of 0.844 (Figure 4A) while our previous map had a resolution of 10Å and a sphericity of 0.676. (We had binned the images to a pixel size of 4.8Å/pixel for our previous structure, but using a smaller pixel size did not improve the quality of the map). Moreover, while a comparable number of particles contributed to both structures (51,119 particles for the older structure and 61,654 particles for the current one), the current structure required much less data processing. To generate our previous map, we performed extensive 2D and 3D classification to remove the preferred views from a dataset of 800,000 particles selected after the initial round of 2D classification. In contrast, for the current structure we selected 350,000 particles after a single round of 2D classification, and after only one additional round of 3D classification (Figure S2) we identified the homogeneous subset of 61,654 particles that yielded an isotropic 3.1Å-resolution map. In the present case, we did not have to remove overrepresented views from the dataset and the improvement in resolution was mainly due to a more uniform Euler angle distribution of the particles (Figure 4B)

**Figure 4:**
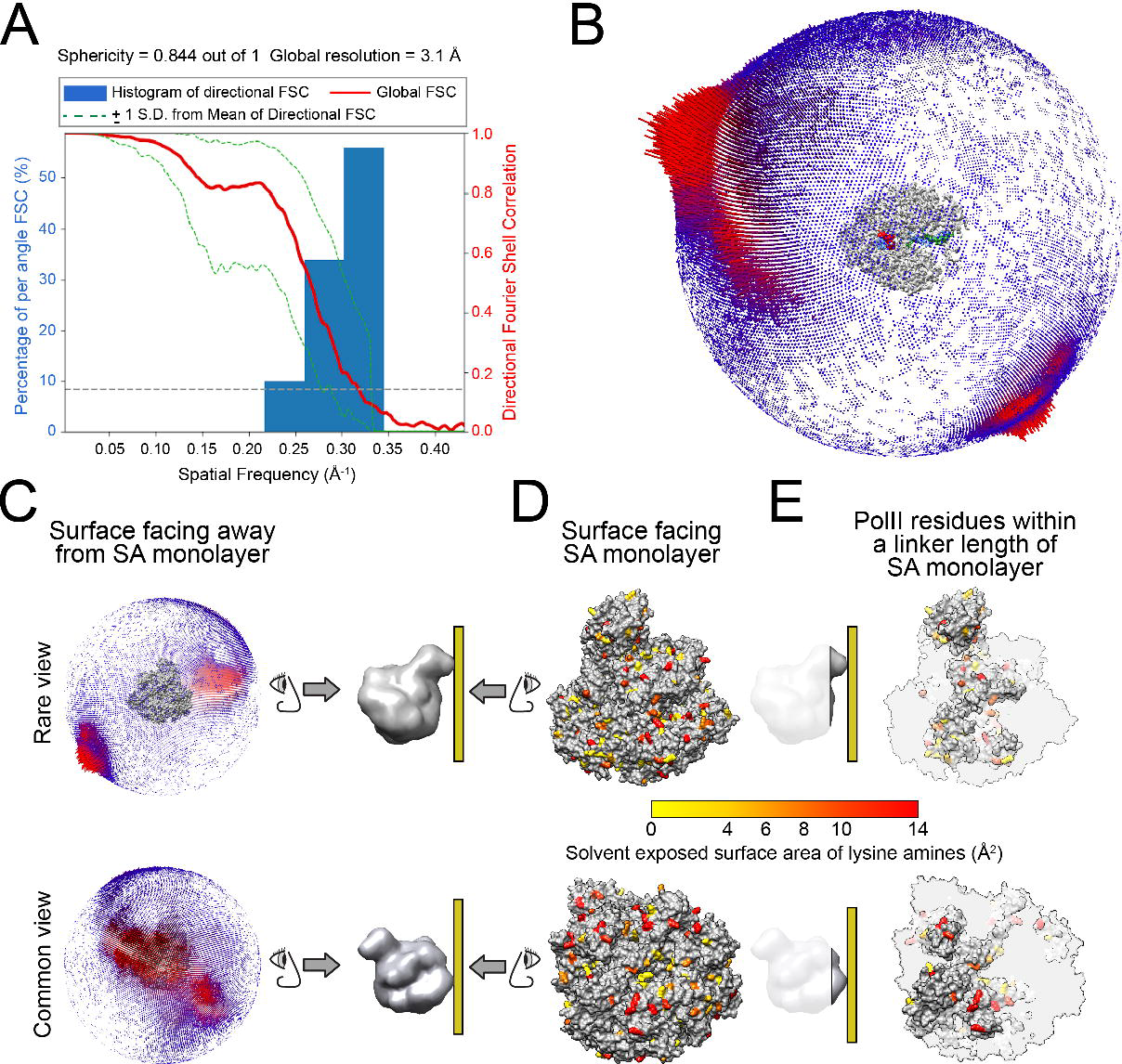
The accessibility of surface-exposed lysines correlates with particle orientation. **(A)** 3D FSC plot for the Pol II EC(CPD) reconstruction. **(B)** Euler angle distribution of the particles contributing to the reconstruction. **(C-E)** Solvent exposed surface area (SASA) of the lysine primary amines available for biotinylation. We used the atomic model of a previously published Pol II elongation complex (PDB ID: 1Y77) to represent the Pol II EC(CPD) in these panels. **(C)** A rare (top) and a common (bottom) view of Pol II EC(CPD) are shown along with the corresponding plots of Euler angle distribution. **(D)** This panel shows the surfaces of Pol II that must face the SA layer to lead to the views shown in (C). The primary amines of lysines are color-coded according to their solvent accessible surface area (SASA), with the least exposed amines shown in yellow and the most exposed ones in red. **(E)** This panel shows the same views shown in (D) but with a cutting plane adjusted such that only lysines whose biotin moieties would be capable of interacting with the streptavidin monolayer are shown. The cutting plane was set to ~29Å from the lysine’s amine, which is the length of a fully stretched biotinylation reagent molecule. The orientations and the cutting planes of the Pol II EC with respect the streptavidin monolayer (ochre rectangle) are shown next to each of the views in (D) and (E).

Despite the lower anisotropy of the map reported here (Figure 4A,B), certain views were still dominant (Figure 4B). One possible explanation for this observation could be the non-uniform distribution of surface-exposed lysines, which might in turn result in an uneven distribution of biotinylated sites on the Pol II EC(CPD). Furthermore, given the finite length of the biotin+linker moiety, some biotinylated lysines may be found in cavities where they cannot reach the streptavidin monolayer crystal. To test whether these two factors could explain the uneven distribution of views we observed, we calculated the solvent accessible surface area (SASA) of each lysine residue’s primary amine. Indeed, we found that those surfaces of Pol II that face the streptavidin crystal in the preferred views contain a larger number of lysine residues with high SASA relative to those representing rare views (Figures 4C and D). This correlation is even more pronounced when we only consider those residues whose biotin moiety would be physically capable of reaching the streptavidin monolayer (Figure 4C and E). For instance, the surface of Pol II corresponding to the rare view shown in Figure 4C (top panel) has ~19 surface exposed lysines with an average SASA of 7Å^2^ (Figure 4E, top) while the surface for the common view in Figure 4C (bottom panel) has ~31 surface exposed lysines with an average SASA of 11.5Å^2^ (Figure 4E, bottom).

### The CPD lesion at the active site induces Pol II backtracking but does not distort the downstream DNA

The structure presented here is largely in agreement with a crystallographic structure of Pol II EC(CPD) (PDB ID: 2JA6) reported by Brueckner et al. with an overall R.M.S.D of 1.1Å (Figure 5A) (Brueckner et al., 2007). As previously reported, we found that the CPD lesion in our structure occupied a position one nucleotide downstream of its expected location (Figure 3F). We designed our scaffold for the CPD lesion to occupy positions +1 and +2, with +1 being the position of the templating base (Figure 3F top panel). However, in our structure we found that Pol II had backtracked by one nucleotide, placing the CPD lesion at positions +2 and +3, downstream of the bridge helix (Figures 3F bottom panel and G). The backtracking leads to a loss of base pairing between the +1 dTMP and the 3’ terminal AMP of the RNA transcript (Figure 3F, bottom panel and G). In fact, we could only detect very fragmented density for the AMP.

**Figure 5:**
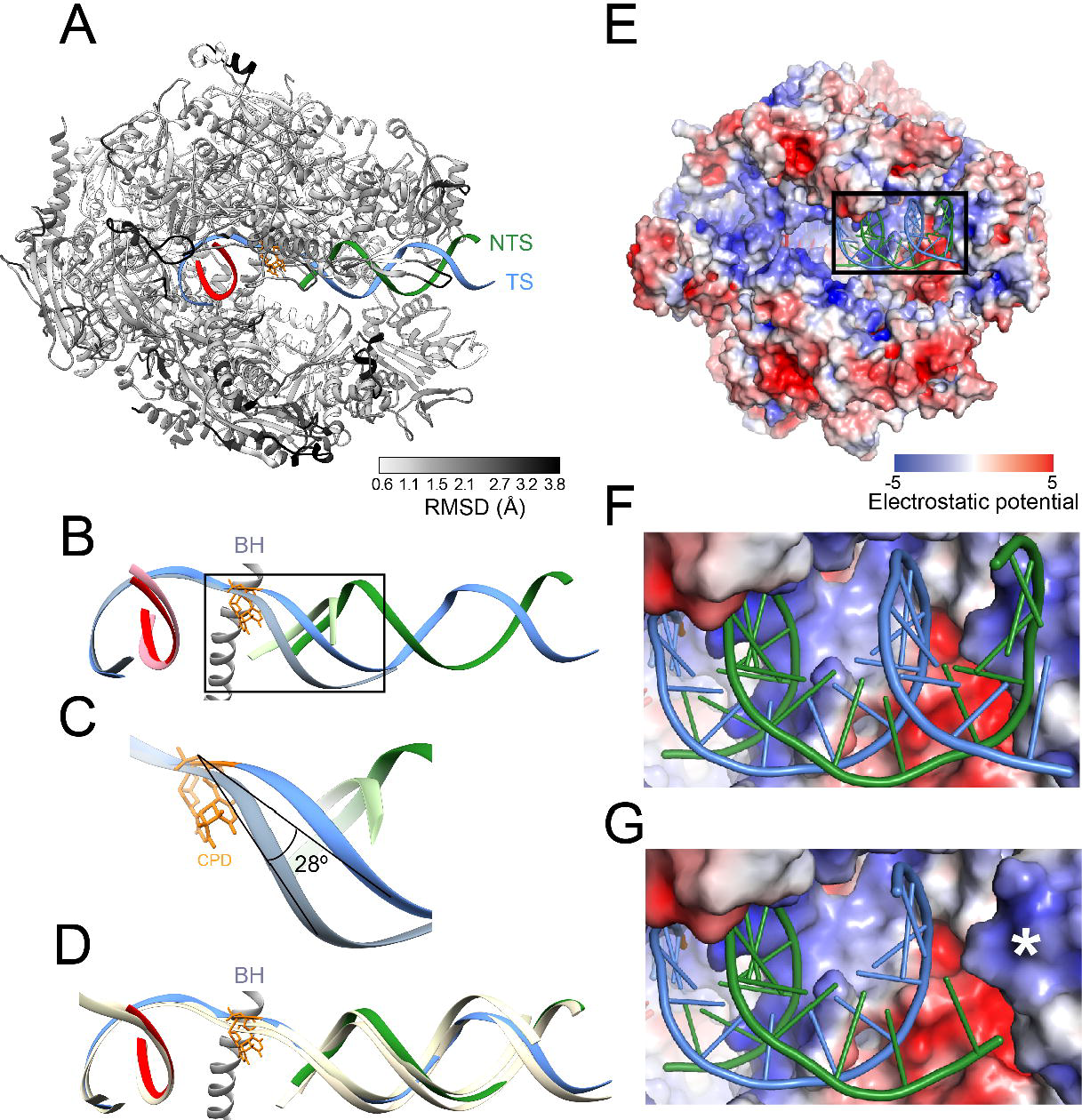
The downstream DNA adopts a canonical orientation despite the local distortion induced by the CPD lesion. **(A)** Comparison between the structure of Pol II EC(CPD) presented here and a previously published structure of Pol II EC(CPD) solved by X-ray crystallography (PDB ID: 2JA6). The root-mean-square deviation (r.m.s.d.) between the Pol II backbones is represented by shades of grey on our structure. The transcription scaffold is colored following the scheme described in Figure 3. **(B)** Comparison between the transcription scaffolds of the two Pol II elongation complexes with CPD-containing DNA templates. The structures were aligned using the protein components; the panel shows the resulting positions of the transcription scaffolds. Our structure was color-coded as in Figure 3. For 2JA6, the non-template strand is in light green, the template strand in light steel blue and the nascent RNA in pink). The bridge helix (BH) and the CPD lesion from our structure are shown in grey and orange, respectively. **(C)** Close-up of the region highlighted in B (black square). The BH was omitted for clarity. The angle between the template strands in 2JA6 and our structure is shown. **(D)** Comparison between the transcription scaffold of Pol II EC(CPD) from this study and those of two elongation complexes with undamaged DNA templates. As in (B), the structures were aligned using the protein components and the panel shows the resulting positions of the transcription scaffolds. The Pol II EC(CPD) from this study is colored as in (B); The structures of the two elongation complexes with undamaged DNA (5VVS and 5FLM) are shown in yellow. The bridge helix (BH) and the CPD lesion from our structure are shown in grey and orange, respectively. **(E)** Interaction between Pol II and the transcription scaffold. Pol II is shown in surface representation and color-coded according to its electrostatic potential, reported in units of K_B_Te_c_^−1^, where K_B_ is the Boltzmann constant, T is the temperature of calculation (310 K) and e_c_ is the charge of an electron. The transcription scaffold is shown in stick representation. **(F)** Close-up of the region highlighted in (E) (black square). **(G)** Same view as in (F) but four bases have been computationally removed from the downstream end of both the TS and NTS to mimic the length of the downstream scaffold in 2JA6. The asterisk highlights a positively charged patch that cannot be reached by the shortened scaffold.

A major difference between our structure and that from Brueckner et al. is in the downstream DNAs, where their paths diverge by ~28° (Figure 5B and C). Unlike Brueckner et al.’s structure, ours matches well those of other Pol II EC structures containing undamaged transcription scaffolds (Figure 5D), despite the well-documented local deformation induced by the CPD lesion (Park et al., 2002). Given that the downstream DNA used by Brueckner et al. was 4 base-pairs shorter than ours, it is possible that it was unable to interact with Pol II in ways that would otherwise compensate for the CPD-induced distortions and maintain its canonical location. In fact, there is an interaction in our structure between the downstream DNA and a region of positive charge in the clamp domain distal from the active site (Figure 5E and F); this interaction could not take place with the shorter downstream DNA used by Brueckner et al. (Figure 5G).

## Conclusions

The 3.1Å structure of a *Saccharomyces cerevisiae* RNA polymerase II elongation complex stalled at a CPD lesion (Pol II EC(CPD)) presented here shows that streptavidin monolayer affinity grids (“streptavidin grids”) can be used to overcome the preferred orientation problem and solve high resolution cryo-EM structures of biological macromolecules. Our previous structure of Pol II EC(CPD), obtained using holey carbon grids, was severely limited in resolution due to the particles adopting strongly preferred orientations(Xu et al., 2017). We could not remove this bias by using grids coated with a graphene oxide support layer or by adding BS3, a cross-linker additive that has helped overcome the preferred orientation problem for certain Pol II transcription complexes (Bernecky et al., 2016; Ehara et al., 2017) (data not shown). In contrast, we obtained the 3.1Å structure presented here from the first streptavidin grid we screened.

We found the streptavidin grids to be very robust in terms of mechanical stability and grid handling procedures and suitable for routine cryo-EM experiments. Moreover, in our hands the random biotinylation of Pol II EC(CPD) did not disrupt the stability of the complex. An additional advantage of this method is its ability to concentrate the sample on the cryo-EM grid, making it particularly attractive for low abundance samples.

Consistent with a previous report, we found that for a biotinylated sample applied to a streptavidin grid the particle orientation is not uniformly distributed(Han et al., 2016). Our results suggest that the particle distribution positively correlates with the distribution of surface exposed lysines (the amino acid residues to which the biotin moieties are attached) and with the ability of biotinylated lysines to reach the streptavidin surface. For known structures, calculating the solvent exposed surface area (SASA) of the lysines might provide an indication of whether the sample will adopt orientations on a streptavidin grid that were absent from data collected using other approaches. To this end we have added a module to Rosetta (will be available from Rosetta 2019.10 onwards) for calculating lysine SASA. The user can perform this calculation as long as some model of the complex under consideration is available.

Compared to the currently available cryo-EM sample preparation approaches for preventing a protein from interacting with the air-water interface, we find the streptavidin grids to be a robust, efficient and reproducible method that can be used with different samples with minimal optimization. This method should help in speeding up high resolution structure determination by overcoming the detrimental effects of the air-water interface.

## Acknowledgements

We thank Robert M Glaeser (UC Berkeley) for his advice and support throughout this project. We also thank the UC San Diego Cryo-EM Facility and the UC San Diego Physics Computing Group. This work was partially supported by NIH grants 2R01GM092895 and 2R01GM107214 (A.E.L); R01GM102362 (D.W.); R01GM123089 (F.D.), and P01GM051487 (Robert M. Glaeser).

## Data accession

The locally filtered cryo-EM map of Pol II EC(CPD) has been deposited in the Electron Microscopy Data Bank (EMDB) with the accession number of EMD-0633. The atomic model corresponding to this map has been deposited in the Protein Data Bank (PDB) with the accession number 6O6C.

---

**Figure S1:**
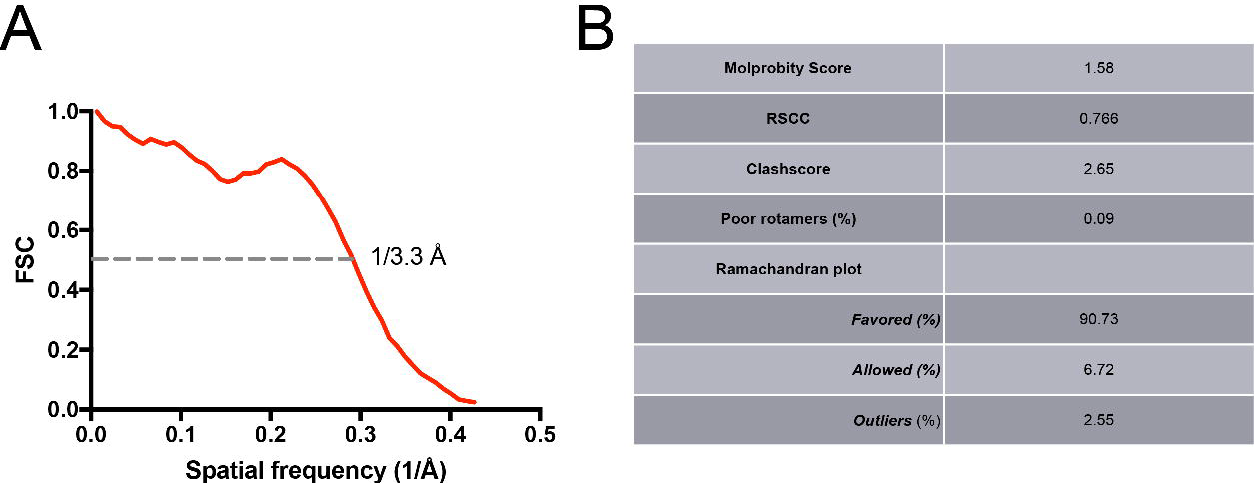
Validation of the atomic model of the Pol II EC(CPD) complex. **(A)** FSC between the atomic model and the cryo-EM map of Pol II EC(CPD) with the spatial frequency at 0.5 FSC indicated. (B) Table summarizing the geometry statistics of the model.

**Figure S2:**
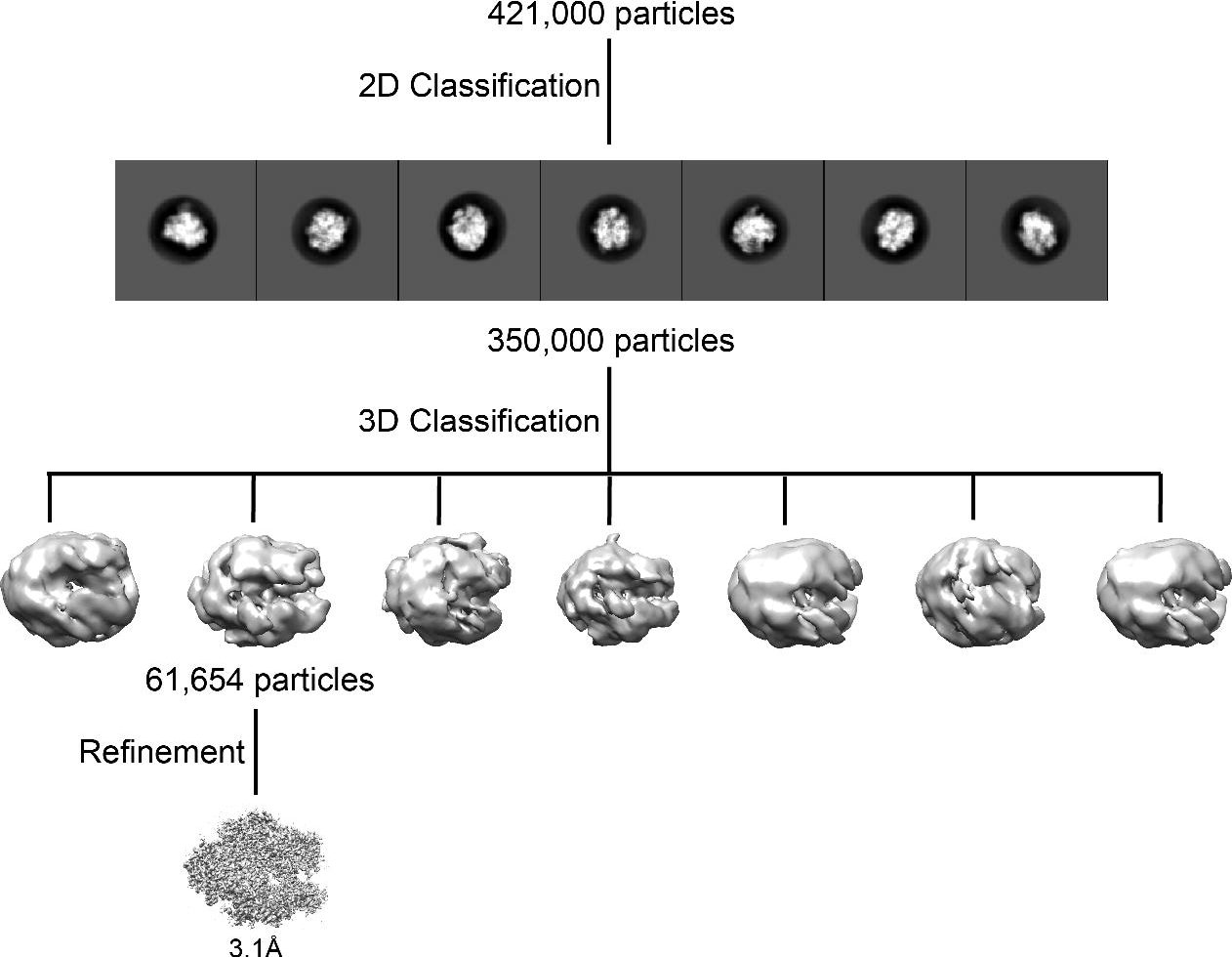
2D and 3D classification strategy used to select the particle subset for the reconstruction shown in Figure 3.

